# The lost world of Cuatro Cienegas Basin, a relictual bacterial niche in a desert oasis

**DOI:** 10.1101/311381

**Authors:** V. Souza, A. Moreno-Letelier, M. Travisano, L. D. Alcaraz, G. Olmedo-Alvarez, L. E. Eguiarte

**Affiliations:** Jardín Botánico, Instituto de Biología Universidad Nacional Autónoma de México, Coyoacán 04510 Mexico City, Mexico.; Department of Ecology, Evolution and Behavior, University of Minnesota, Saint Paul, MN 55108; Laboratorio Nacional de la Sustentabilidad, Instituto de Ecología, Universidad Nacional Autónoma de México, Coyoacán 04510 Mexico City, Mexico.; Laboratorio de Biología Molecular y Ecología Microbiana, Departamento de Ingeniería Genética, Unidad Irapuato Centro de Investigación y Estudios Avanzados, IPN, México.

## Abstract

Barriers to immigration can lead to localized adaptive radiations and increased endemism. We propose that extreme oligotrophy can be a strong barrier to immigration over geological timescales, and facilitate the evolution of diverse and coevolved microbial communities. We show here that the endangered oasis of Cuatro Ciénegas Basin can be a model for a *lost world*, where the ancient niche of extreme oligotrophy favoured survival of ancestral microorganisms that persisted due to environmental stability and low extinction rates, generating a diverse and unique bacteria diversity. Diversification/extinction rates in *Bacillus* showed several CCB endemic clades that diverged from the rest of *Bacillus* spp. in different times of the Paleozoic and Mesozoic, in contrast to more recent *Bacillus, Clostridium* and *Bacteroidetes* lineages. CCB conservation is vital to the understanding of early evolutionary and ecological processes.

## Main

A “lost world” is both a poetic metaphor and a scientific pursuit; in both cases, it pertains the conservation or recreation of the deep past in a particular place. Scientists have looked for analogues of such world in environments possessing living microbial mats and stromatolites, since these structures were dominant for billions of years in the Proterozoic^1^. Nevertheless, in most cases, these communities represent more a physical metaphor of the past than an actually lost world, since they harbour mostly derived microbial lineages^2,3^. The exception to these observations seems to be the abundant and morphological diverse stromatolites and microbial mats from the endangered oasis of Cuatro Ciénegas Basin (CCB) in Northern Mexico. In this extremely diverse wetland^4^, the recycling of the deep aquifer by magmatic heat replicates many conditions of ancient oceans^5^ including its extreme oligotrophy with phosphorous as low as 0.5 mM^6^. Spring water is also low in oxygen and rich in sulphur minerals^5,7^. These conditions represent niche variables that can explain the survival of marine as well as hydrothermal vent associated sulphur microbes^8^. Viral metagenomics confirm these observations, as these studies have shown substantial divergence from known sites and a similitude with marine habitats^9^. All these data have been hinting at a lost world scenario, since CCB, now it at the centre of the North American continent, was at its shore 35 Ma ago^8^.

If extremely low phosphorous levels were the initial conditions where these microbes evolved, bacteria have been, since their beginning, under strong selective pressure to adapt to these environmental conditions, generating parallel new ways to obtain the element that is rare. Nutrient conditions create an extreme stoichiometric bias in the N:P ratio (as high as 122:1 in our study site, Churince) at the community level ^10^, but also an extreme imbalance at the bacterial cellular level (N:P 965:1 in a CCB *Bacillus cereus* group ^11^). Extreme oligotrophy, as well as rich sulphur conditions are a specific niche characteristic of the Precambrian ocean, that ended abruptly in the Phanerozoic Eon 542 Ma^12^. Moreover, using conservative time frames based on geological events, molecular clock studies have suggested that some strains of cultivable cyanobacteria^13^ as well as of *Bacillus*^14^ from CCB diverged many millions of years ago from their close relatives.

Hence, we propose that CCB is a microbial lost world, not just as a poetic metaphor, but as a true geographical site; a nutrient-deprived multidimensional niche isolated from the human environment by its peculiar conditions, as calcium carbonate rocks work as a buffer between the human activities and the turquoise blue ponds^5^.

What would make a lost world more than a metaphor? If we compare different models of diversification (Fig. 1), we can observe that for cosmopolitan bacteria, such as *Escherichia coli* or *Bacillus subtilis*, the phrase “everything is everywhere and the environmental selects” could apply, since these microbes have an enormous population size and considerable migration rates (1a). A second model is isolation by distance, a pattern similar to the one observed for most macro-organisms. This isolation has been observed for the thermophile Crenarchaeota *Sulfolobus islandicus*^15^(Fig. 1b). The third model is that of isolation as in an island-like migration (Fig. 1c), and this pattern also occurs in microbes, as is the case for those in the lakes in the Pyrenees, explained by the island-like nature of each lake^16^. Here we suggest that there is a 4^th^ model, the lost world model (Fig. 1d), where a particular site is maintained as a “bubble-like” niche, a place where a community survived in relictual conditions. In this case, we would observe phylogenies where a few isolated microbes would exhibit long phylogenetic branches, and even complete lineages would appear separated and diversified from others in parallel bouts.

**Fig. 1.**
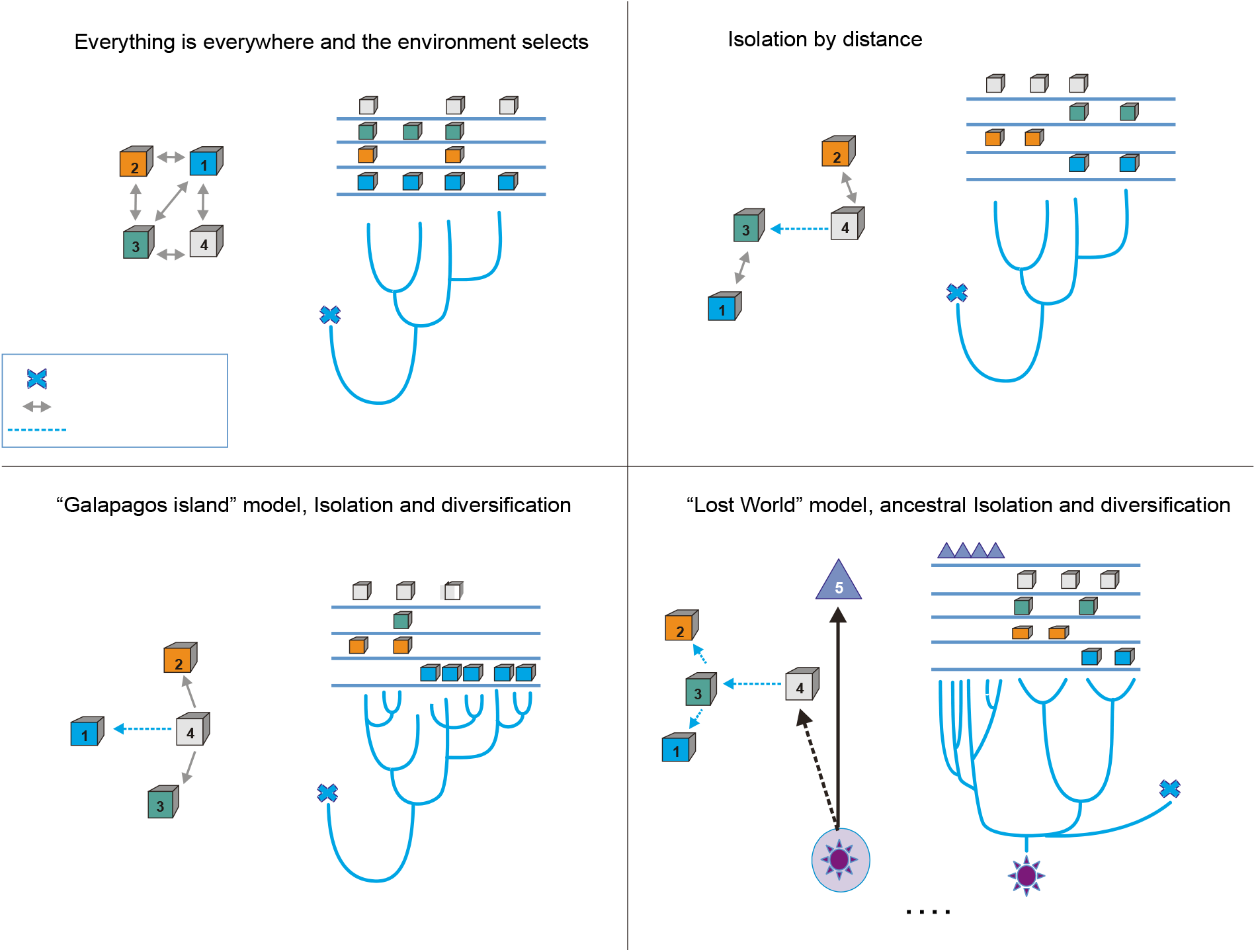
Conceptual frame-work of species diversification. A) “Everything is everywhere, and the environment selects” model implies that free migration is only restricted by environmental filtering. Hence, sites 1-4 have the same probability of migration, but some sites, such as 1 are better than others. In this case, the phylogenetic tree does not reflect the geographic structure; this is common in many cosmopolitan lineages of bacteria and fungi. B) Model of isolation by distance, this is what occurs in most plants and animal phyla, sites that are closer (1 and 3 or 2 and 4) are more likely to present migration events than sites that far apart, some rare events of migration are allowed (as between 4 and 3). In this case, the branches of the tree reflect the geographic structure. C) Island model implies that rare events of migration from the source (4) to an island (1). The phylogentic tree reflects adaptive radiation due to isolation. D) “Lost world model” of ancestral isolation and diversification implies that lineages that were extinct in other places have remained as relictual niches persist in a new site (5). In this case, the ancestral lineages have very long branches that show their ancestral diversification from common ancestors.

In order to distinguish between these four models in our system of study (CCB), we will first describe the microbial diversity at a small scale (ca. 1 km^2^) and then zoom-in even further into particular cultivated genera within the Gram positive. We studied aerobic (*Bacillus*) and anaerobic (Clostridiales and Bacteroidetes) lineages, to have a sample large enough to evaluate how much the CCB lineages diverged from those in the rest of the world.

Using NextGen 16S rRNA gene tags, we surveyed the microbial biodiversity in the Churince, a closed hydrological system^11^ (and the most endangered site within CCB). We observed a vast diversity (Figure 2) on a small scale (less than 1 km^2^), in particular when we compared such diversity with other 343 16S rRNA studies and microbiomes. Those studies comprised several contrasting environments: human, plant-associated, soil, sediments, biofilm, marine biofilms and extreme environments like Yellowstone hot springs, Guerrero Negro salt flats, and Antarctica soils, all available in public databases (Table S1). Based on standard DNA similitude metrics, an operational taxonomic unit (OTU) is defined in general as 97% identity at the 16S rRNA gene.

**Fig. 2.**
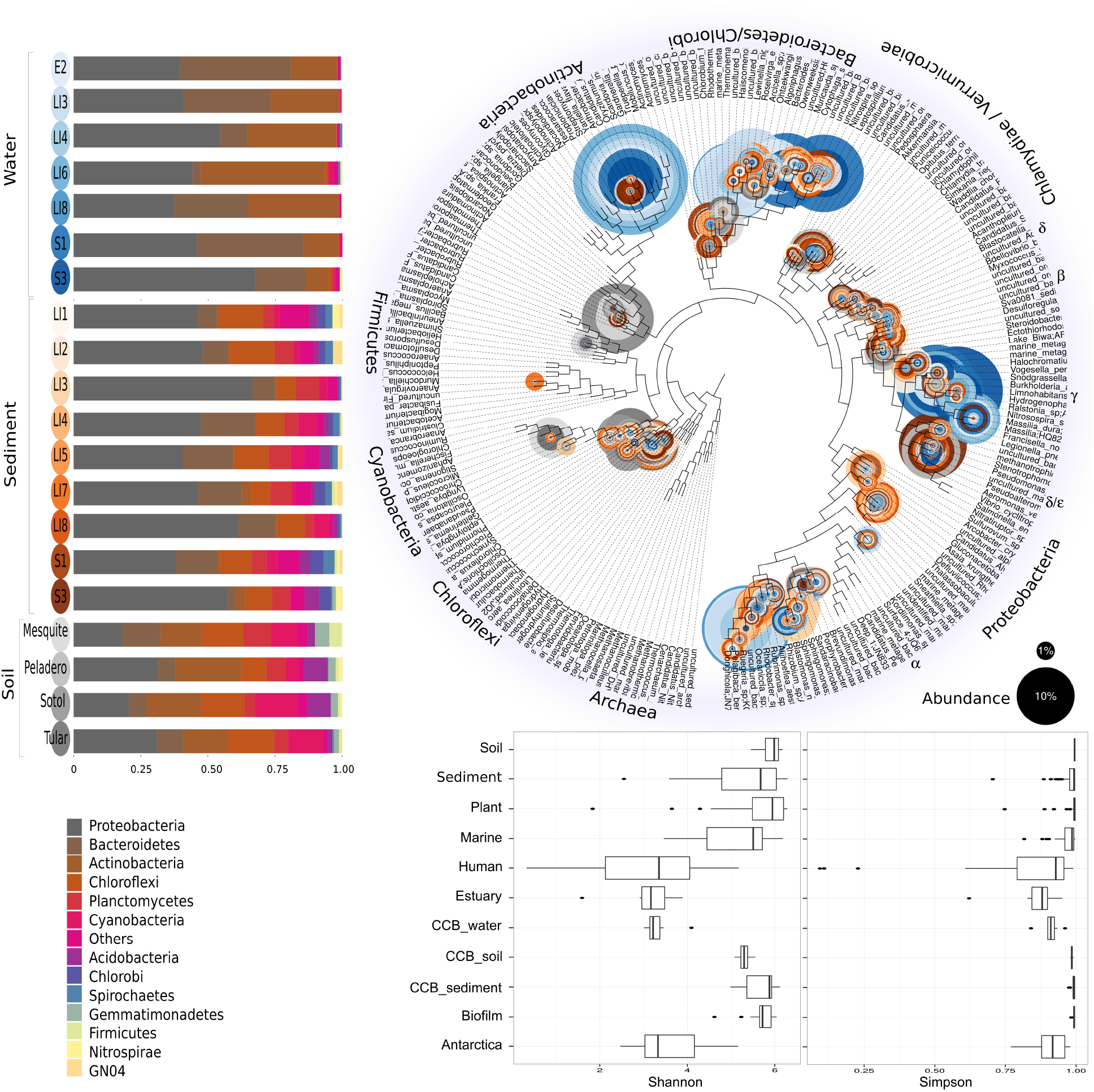
Overall prokaryotic diversity in Churince. Major phyla abundances in CCB is depicted in bar plots, only the most abundant phyla are shown, but there are 60 phyla present in CCB which are roughly 66.28% of known prokaryotic phyla, in a single location few meters away. Proteobacteria is the most abundant phylum, followed by Bacteroidetes, and Actinobacteria. Some phyla like Planctomycetes, Cyanobacteria, Acidobacteria, Chlorobi, and Firmicutes are more abundant in the sediment and soil-associated samples than in water columns. In the right panel, a phylogenetic placement shows the names and abundances of the most represented genera within CCB samples. Each CCB sample is colour-coded according to its origin: blue for water; brown for sediments; and grey for soils. Note that some genera of *Actinobacteria, Bacteroidetes/Chlorobi*, a-Proteobacteria, and γ-Proteobacteria, are only in water samples, while sediment and soil samples are represented all across the tree. In the lower right panel, Shannon and Simpson diversity indexes for 342 microbiomes, and metagenomes retrieved from public databases were calculated and compared to diversity values of CCB. The most diverse environments were soil, aquatic sediments, and plant-associated microbiomes, both of the top scores in Shannon’s and Simpson’s index. CCB is a paradox of nutrient scarcity and quite high diversity, possibly a product of a long-term evolutionary scale, complete food chains, and stable interactions.

The Churince’s total Bacteria and Archaea richness are represented by a total of 5,167 OTUs assigned to samples from the water column, aquatic sediments and soil (Fig. 2a). The assigned OTUs represented 60 different known phyla, three of which were Archaea. To compare with the other environments, we used the individual OTUs and then computed two alpha diversity indexes: Shannon, and Simpson (Fig. 2b).

The most diverse environments according to Shannon’s diversity index are the aquatic sediments of different sites in the world; accordingly, the Churince’s sediment had the highest Shannon value of our dataset. This was also supported by the Simpson’s index, showing that water column < soil < sediment (Fig. 2b). Using both Shannon’s and Simpson’s diversity indexes, Churince showed a very high microbial diversity, even when compared with other microbial diversity hotspots, such as Pearl river in China, or Guerrero Negro in Mexico (Table S1).

To test for the lost world clade diversification, we zoomed-in into the diversity of CCB cultivated *Bacillus*, Clostridiales and Bacteroidetes. From our collection of approximately 2,500 cultivated *Bacillus* spp. from CCB, 265 unique sequences were obtained, selected at 97% identity, a very conservative estimate for *Bacillus*. In a global tree (Fig. 3), we observed that endemic CCB strains formed many exclusive (only feoun in CCB) lineages when we added them to the 1,019 OTUs reported for *Bacillus* spp. from around the world (Fig. 3a), increasing by nearly 25% the number of known *Bacillus* species.

**Fig. 3.**
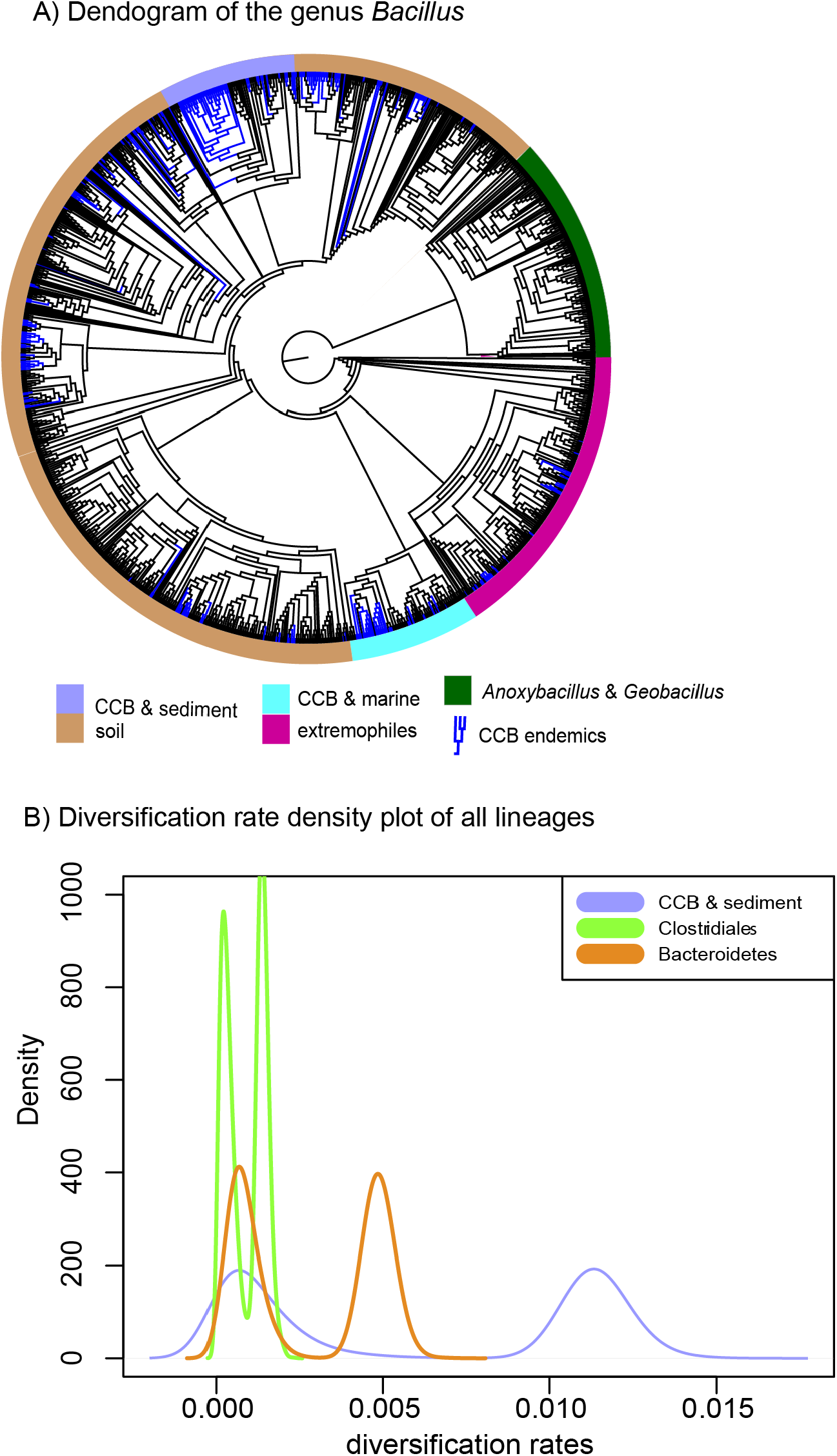
A) Dendrogram of the 1,284 strains of the genus *Bacillus* reconstructed from 16s rRNA. All taxa have a sequence divergence over 97%. Strains found in Cuatro Cienegas Basin (CCB) are denoted in blue. All other strains, including the outgroups, are denoted in light gray. The position of the sediment CCB *Bacillus* and marine CCB *Bacillus* lineages within the genus *Bacillus* are also indicated. The outgroup includes strains of *Geobacillus* and *Anoxybacillus*. B) Density plot of the rates of diversification of Bacteroidetes, Clostridiales and sediment CCB *Bacillus*. All lineages show a bimodal distribution of diversification rates, with sediment CCB *Bacillus* having rates one order of magnitude higher than Bacteroidetes.

Within the *Bacillus* spp. from CCB, we can distinguish two diverse endemic lineages: one from sediment and another one closely related to marine *Bacillus* spp. (Fig. 3). Sediment CCB *Bacillus* spp. are significantly older than the marine related CCB lineages and according to our analysis date back to the Ediacaran, the period that holds evidence of the oldest multicellular organisms (Fig. S3, period after the Cryogenian at the end of the Precambrian). Unlike the sediment lineage, marine related CCB *Bacillus*species did not form a monophyletic group, which suggests independent synchronized origins dating to the late Jurassic (Fig. S4). It seems as if a mixed representation of many lineages entered this “multidimensional niche bubble” simultaneously and diversified according to the new rules of interactions along with the neighbours. The rates of diversification of sediment and marine related CCB lineages suggest a lost world scenario for both (Fig. 1D).

None of the other lineages of *Bacillus* exclusive to CCB have been reported in any other site but they seem to constitute later events of arrival to the “island-like” unique niche and to have diversified locally. Bacteroidetes and Clostridiales strains were fewer and did not form monophyletic groups endemic to CCB (Figure S1), even though the Bacteroidetes had a bout of diversification at the end of the Paleozoic (Figure S2), suggesting also punctuated migrations and local radiations.

Environmental conditions are known barriers to dispersal and the extreme P limiting conditions at CCB could certainly constrain immigration from P demanding populations. This is the case of Patagonia’s isolated and oligotrophic lakes that present unique but low diversity^17^. CCB is not only unique, but it also very diverse, despite its extreme oligotrophy. Our data showed that the small and endangered Churince system in CCB, contains 57 out of the 86 known Bacteria phyla, which is 66.3% of the world recorded bacterial diversity at phyla level (data from 342 analysed microbiomes, Table 2S). This diversity is only comparable to Pearl River in China^18^, where 48 microbial phyla are found. In contrast to CCB, Pearl River is a highly productive environment that receives inputs from multiple sources in the 2,400 km extension of China’s third largest river, while Churince is a small hydrological system fed by a spring that extends 1 km at most.

The local scale makes the bacterial diversity of CCB even more interesting. It is possible that the reason for such diversity is not only due to local adaptation to the environment but to a “Red Queen” evolutionary process, where interactions among species shape diversity, causing an increase in localized adaptation and coevolution^19^. Experiments of competition between strains of *Bacillus* from different sites within Churince showed marked antagonisms against different strains from even sites few meters away, but tolerance and even cooperation between strains from exactly the same site^20^; for instance, strains of endemic lineages of *Bacillus* from Churince require cross feeding and cooperation to obtain even basic amino-acids ^21^. Hence, one potential explanation for the long-term survival of lost world *Bacilli* species in CCB would be their coevolution with the community members that exchanged nutrients with them in their particularly low nutrient niche following a “Black Queen” model ^22^, maybe originally as part of microbial mats or stromatolites in the South-Western shores of Laurentia^23^.

Our data showed that the total known phylogenetic diversity of *Bacillus* spp. increased by ¼ the known species when we included the strains unique to CCB. Moreover, a large part of this unique diversity belongs to early divergent clades. Even more so previously, some CCB *Bacillus* strains^14^, as well as Cyanobacteria^15^, their divergent dates for other linages were dated molecularly towards the late Precambrian. *Bacillus* spp. are capable of forming spores, the ultimate strategy to survive stressful conditions, however, we have shown that the*Bacillus* spp. at CCB, are not merely in the form of dormant spores, but competing actively in the microbial communities^20^.

Bacteroidetes and Clostridium in CCB seem to follow, like some of CCB *Bacillus* spp., an island model with more recent dispersal events, as the short branches in the phylogenies support a more cosmopolitan origin (Figure S2). Similar patterns were found in a study of *Bacillus* spp. from diverse environments in India where cosmopolitan *Bacillus* spp. had short branches to sister species^24^. All our results suggest that extreme oligotrophy, along with community cohesion function like a “semipermeable” barrier to migration, where effective migration is possible, but rare.

Fossil evidence shows that stromatolites were still abundant between the Permian and Triassic boundary, in the site where the Tethys sea opened in the SouthWestern shores of Laurentia^22^ where CCB was located at the onset of the Mesozoic^8^. However, during the massive extinction event that marked the end of the Permian, stromatolites became rare, except on the western shores of the Tethys sea ^22^. Microbial mats and stromatolites can be still found in other sites of the planet, but at CCB, aside from giving testimony to the past, microbial lineages have been safeguarded, bringing evidence for a lost world.

Even though CCB microbial communities have survived for an extended period of time, their particular niche conditions are being destroyed in the Anthropocene. This impact is even more poignant because CCB wetland has shrank 90% in the last 50 years, and its deep aquifer has been devastated by the use of fossil water in local agricultural practices. This deep niche change has already destroyed many of the microbial complex communities in Churince. However, we believe that this change can be reverted if the channels that drain the wetland are closed and the wetland recovers its water cycle. Conservation of the unique niche in CCB and similar sites is paramount for our understanding of the deep past as well as to predict and protect the future of our planet.

## Acknowledgments

This research was supported by funding from WWF-Alianza Carlos Slim, Sep-Ciencia Básica Conacyt grant 238245 to both VS and LEE, and Sep-Ciencia Básica Conacyt grant 220536 to GOA. The paper was written during a sabbatical leave of LEE and VSS in the University of Minnesota in Peter Tiffin and Michael Travisano laboratories, respectively, with support of the program PASPA-DGAPA, UNAM. We thank Laura Espinosa-Asuar for making Figure 1 and Chris Dupont of JCVI for the phylogenetic analysis in Figure 2.

## Methods

### Microbial diversity context

We sampled ten sites during May 2011 in the Churince system of CCB in a 300 m long lagoon plus two more sites in the spring-head, ca. 1 km away. In each sample site we took 50 g of sediment and a gallon of water as well as a sample of both for biogeochemical variables, nutrients and minerals: C, N, P, Ca, Mg, ^10^. We also sampled four types of vegetation from an established gradient and obtained composite soil samples. We extracted DNA from each sample using the same methodology^25^. Metagenomic DNAs were sent to JCVI (San Diego, CA, USA) for 16S rRNA amplicon gene library (341F-926R primers) 454 pyrosequencing (Roche, Brandford, Ct, USA).

A total of 950,000 reads were sequenced; we required a minimum of 50,000 reads per site, with a minimum 500 bp length after Quality Control check. Not all samples produced the same amount of sequences, probably due to the natural low yield of DNA extraction in CCB water and sediments. Nevertheless, even at 97%, diversity is high, encompassing all the know phyla of Bacteria but a very low diversity and abundance of Archaea and mostly none of the cosmopolitan human related microbial taxa.

The 16S rRNA gene analysis was done as previously reported^26,27^. Briefly sequencing quality was processed and filtered using FASTQ and Fastx-toolkit, we filtered out any sequence with Phred < 30, length < 500 bp. Operational taxonomic units (OTUs) were clustered using cd-hit-est ^28^ with a 97% identity threshold cut-off. The OTUs were parsed into QIIME pipeline and the taxonomic assignments were done against Greengenes DB (v 13.8^29^). Chimeras were removed after taxonomic assignments and detected by ChimeraSlayer ^30^. Data management, diversity statistic, and plots were done using R phyloseq package ^31^and ggplot2 and RColorBreweer R libraries. Pplacer was used to place the diversity into a reference tree (Figure 2) ^10^. We are using diversity indexes, rather than OTU comparisons because of differences in sequencing technologies, primers used for 16S rRNA gene, coverage depth, and other factors that could affect an overall OTU comparison among different studies.

Compared datasets (342) were retrieved from public available databases like NCBI’s SRA, MG-RAST, and HMP (Human microbiome project) websites. Detailed information about accessions used is available as supplementary material.

### *Bacillus* tree

The sequence identity clustering of all 16S rRNA gene sequences from the genus *Bacillus* spp. and sister genera *Anoxybacillus* and *Geobacillus* were retrieved from online databases, 1019 of them at 97% sequence identity, plus 648 sequences of cultivated *Bacillus* spp. from CCB (Genebank numbers in Table S2) selected with the same criterion out of more than 2,500 cultivates strains; sequence clustering was done with cd-Hit (www.cd-hit.org^28^).

These sequences were further aligned with the 16s rRNA sequences from CCB with the MUSCLE plugin in Geneious^32^. Neighbour-joining trees were constructed using genetic distances, with the ape and seqinr R package. The sequences were also aligned with the 16s rRNA sequences from CCB with the MUSCLE plugin in Geneious 5.4.6.^32^ In order to have a control, a subset of OTUs from Clostridiales (n=131; 18 unique from CCB) and Bacteroidetes (n=189; 12 unique from CCB, Genebank numbers in Table S2) was then used to construct Bayesian phylogenies including CCB cultivated anaerobic strains while for *Bacillus*, we only focused on a mayor clade which had a better representation from CCB samples, henceforth called sediment Bacillus (n=311; Genebank numbers in Table S2).

Phylogenies were reconstructed using BEAST v. 1.8.2^33^, with a Birth-Death speciation model, relaxed lognormal clock models and the following substitution models Bacteroidetes (HKY+I+G), Clostridiales (HKY+I+G), and both Bacillus clades (GTR+I+G). All substitution models were chosen using a Bayesian Information Criterion on likelihoods estimated with jModeltest 2.1.7^34^. Three separate runs were performed for each dataset of 50 million chains each and then combined using LogCombiner v1.8.2. Parameter convergence was evaluated using Tracer v. 1.6.0. Ultrametric trees were obtained with relative node ages, which were later scaled to produce ultrametric trees with absolute ages to be used in the diversification rate analyses using the R package phytools^35^.

The heights for each lineage were obtained from the literature. The tree height of Clostridiales was set at 3,500 Ma^14^, for Bacteroidetes was set to 2,500 Ma^36^. The node heights of the sediment *Bacillus* lineage were obtained by estimating the divergence dates within the genus *Bacillus* using a smaller phylogenetic sampling. The analysis was conducted using BEAST v. 1.8.2^33^ and the calibration points were set at the divergence of the genus *Bacillus* from *Geobacillus* at a conservative 1,144.7 Ma (sd= 164)^14^ following the great oxidation event, set to a normal distribution. Another calibration point was set in the diversification of *Bacillus* at 1047 Ma (sd= 159), also set to a normal distribution, as it is the recommendation when using node ages estimated by molecular dating (see BEAST v.1.8.2 documentation). The clock model was a log normal relaxed clock and the analysis was run in BEAST v. 1.8.2^33^.

Changes in diversification rates were estimated with a Bayesian framework using BAMM 2.5.0^37^. This method estimates the speciation rates, identifies shifts along the phylogeny and estimates the confidence intervals of the various shift configurations detected using a Markov Chain Monte Carlo to explore the universe of candidate models^37^. The analyses were carried out using the scaled ultrametric trees of sediment *Bacillus*, Bacteroidetes and Clostridiales, with 10 million generations for all cases except Clostridiales, which required 20 million generations to reach convergence and adequate effective sampling sizes. Priors were estimated using the function setBAMMpriors implemented by the R package BAMMtools. Results were analysed using BAMMtools on R (R Development Core Team 2017; ^38^) to obtain the best shift configuration, Bayes factors of number of shifts and posterior probabilities of shifts distributions.

Finally, we compared the distribution of rates along the tree of all lineages in order to assess the relative diversification rate differences in all lineages.

## Supplementary material

### Supplementary figures

**Fig. S1.-.**
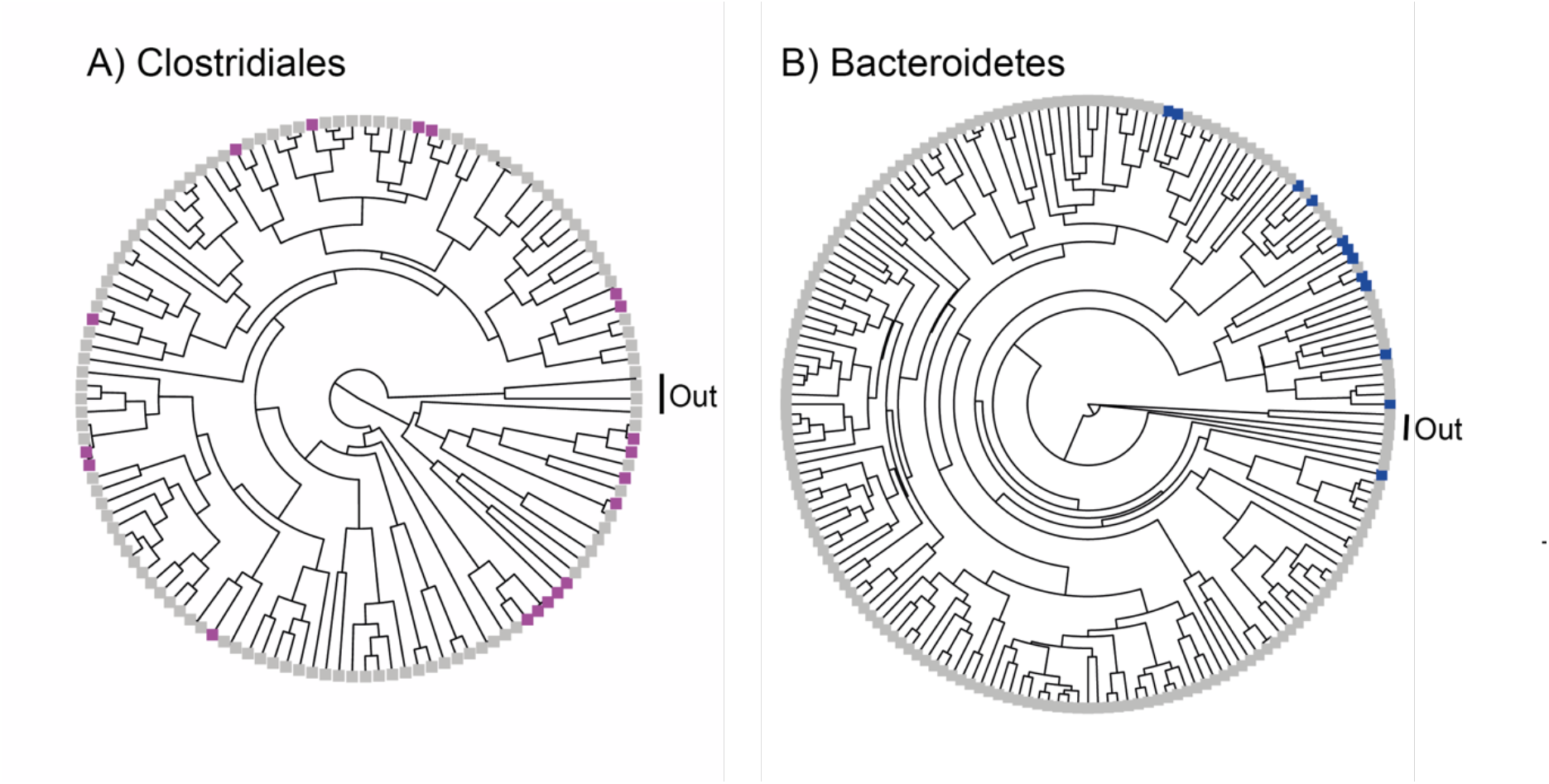
Bayesian phylogenetic trees of the order Clostridiales and the phylum Bacteroidetes reconstructed with 16s rRNA. All taxa have a sequence divergence of 97%. Taxa exclusive to CCB are denoted in magenta for Clostridiales and blue for Bacteroidetes. All other taxa, including outgroups, are in light gray. The outgroup in Clostridiales includes *Thermoanaerobacter* and *Symbiobacterium* and for Bacteroidetes the outgroup includes *Chlorobium* and *Prosthecochloris*.

**Fig. S2.-.**
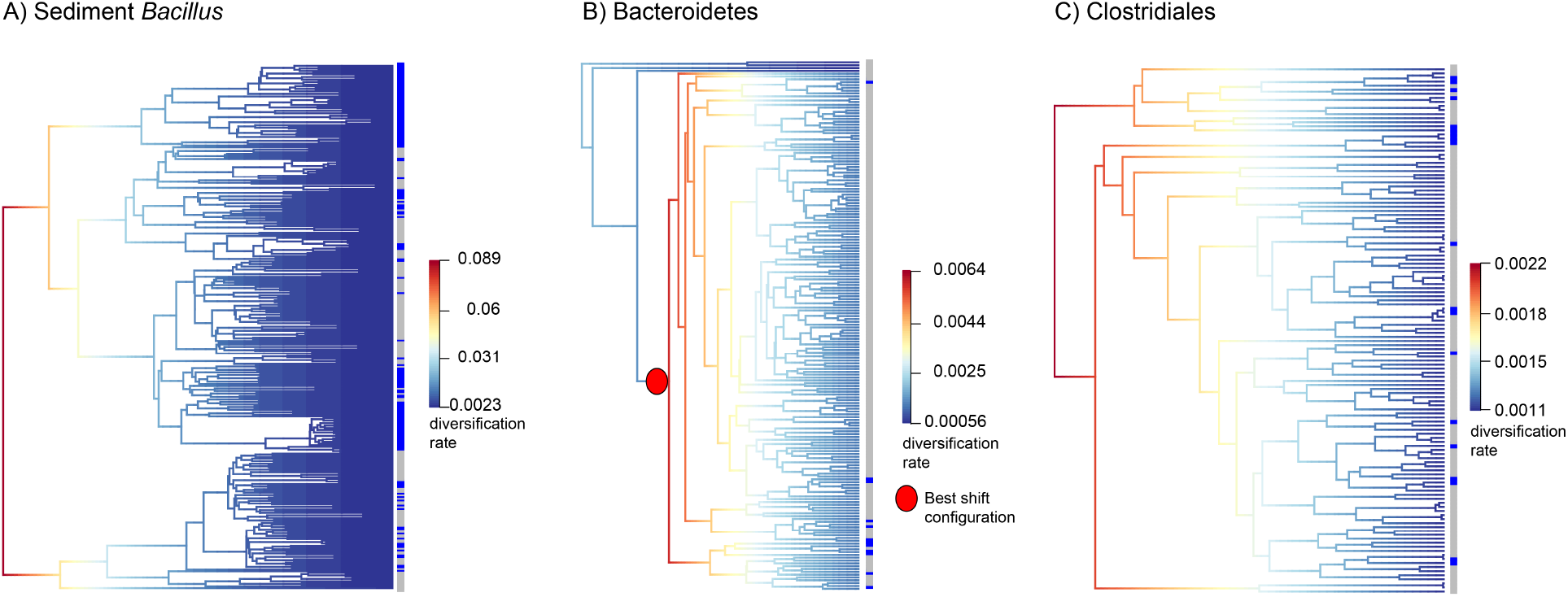
Diversification rates in sediment CCB *Bacillus*, Bacteroidetes and Clostridiales. A diversification rate shift was detected in Bacteroidetes (B).

**Fig. S3.-.**
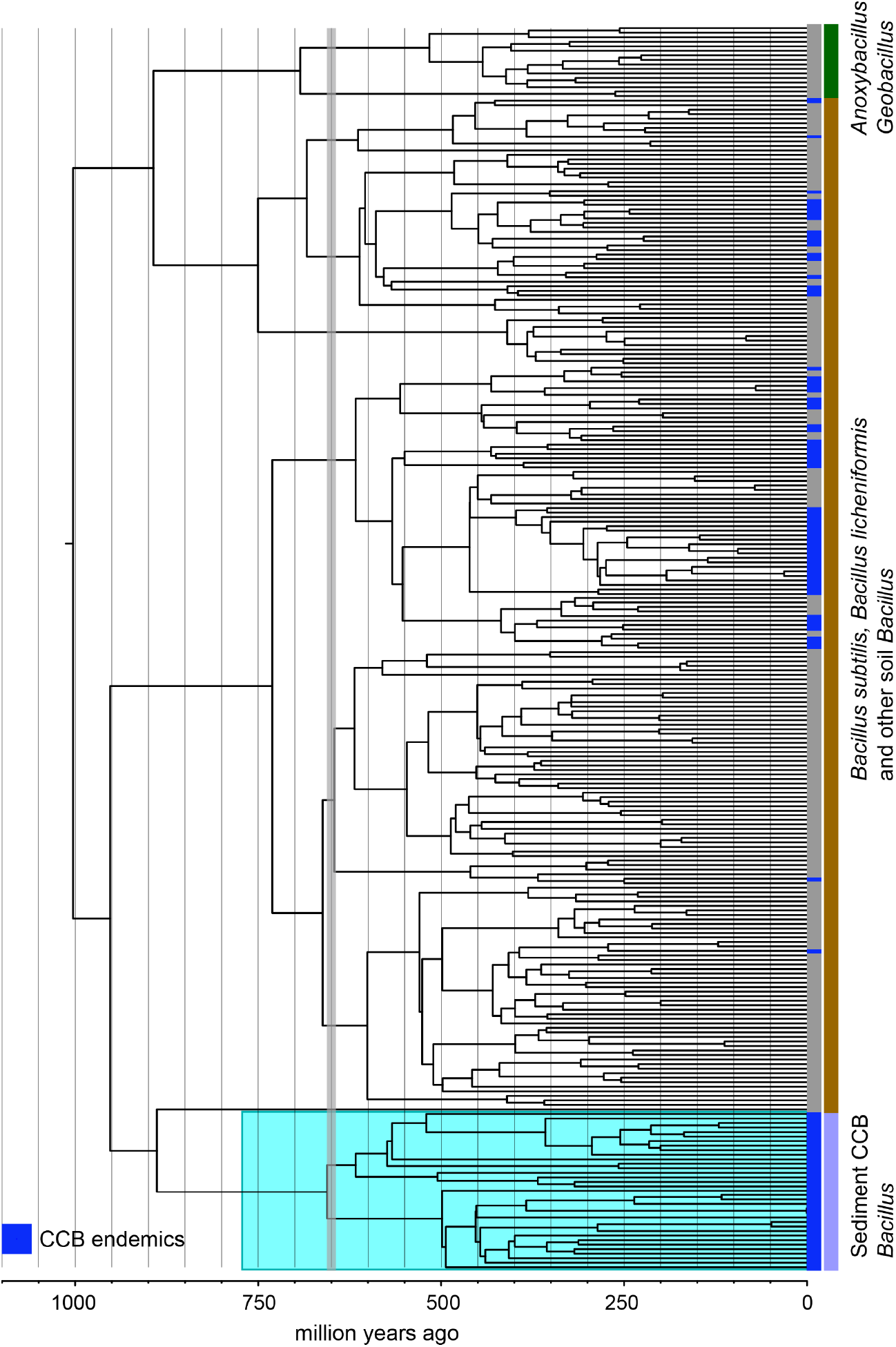
Dated Bayesian phylogeny of soil and sediment *Bacillus*, including the endemic lineage of sediment CCB *Bacillus* (highlighted in cyan). Strains endemic to CCB are denoted in blue. The vertical grey line indicated the date of divergence of sediment CCB *Bacillus* approximately 655 Ma, in the late Precambrian, during the Cryogenian period.

**Fig. S4.-.**
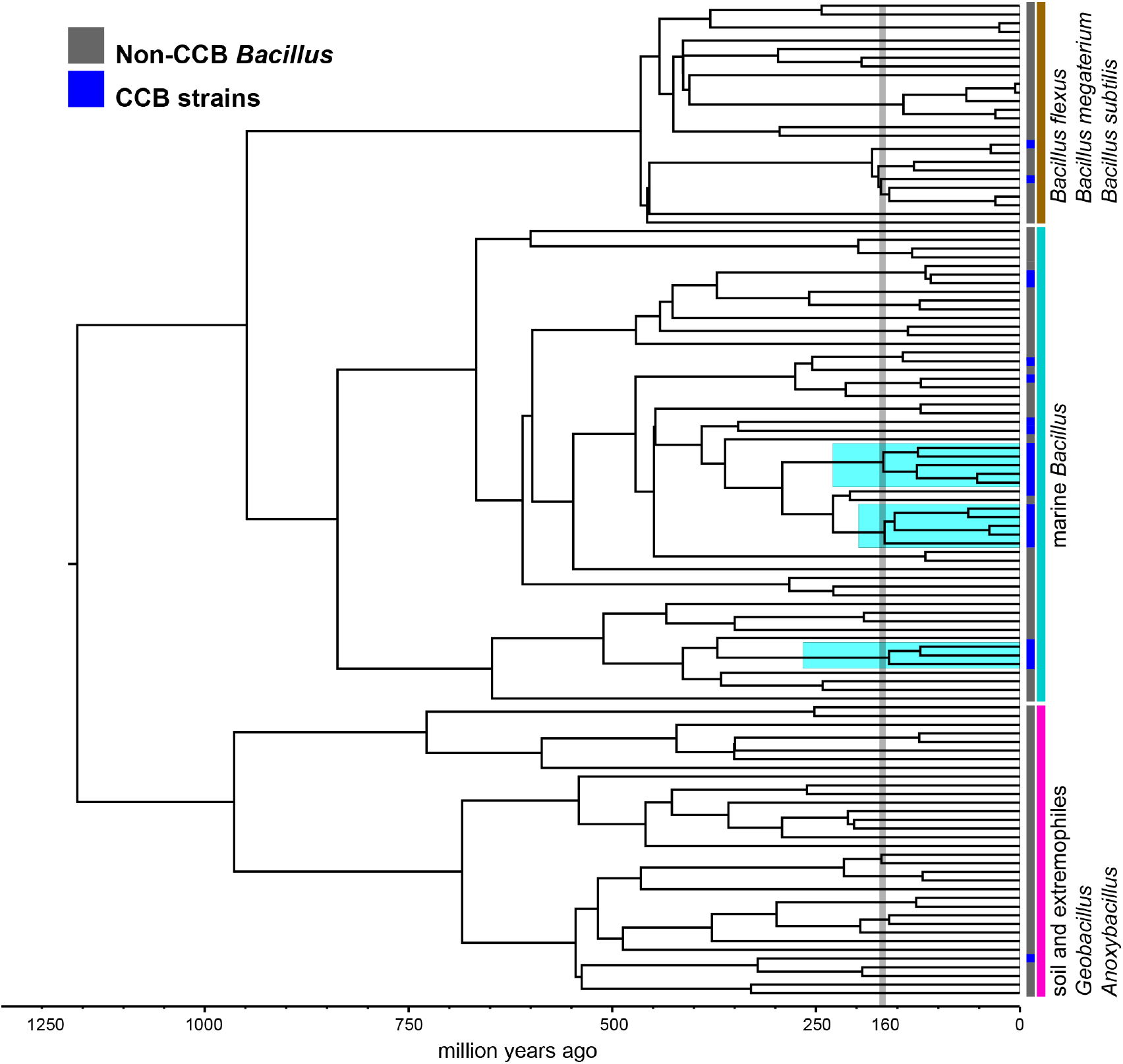
Dated Bayesian phylogeny of marine *Bacillus*, including endemic lineages from CCB (highlighted in cyan). The grey line indicates the divergence time of three independent CCB marine strains at around 160 Ma in the Late Jurassic period.

### Other supplementary material

Table S1.- Churince’s diversity by site, CCB_rest_of_the_world_diversity.zip

Table S2.- Genebank accessions numbers of strains used in this study

